# Pan-cancer screen for mutations in non-coding elements with conservation and cancer specificity reveals correlations with expression and survival

**DOI:** 10.1101/182642

**Authors:** Henrik Hornshøj, Morten Muhlig Nielsen, Nicholas A. Sinnott-Armstrong, Michał P. Świtnicki, Malene Juul, Tobias Madsen, Richard Sallari, Manolis Kellis, Torben Ørntoft, Asger Hobolth, Jakob Skou Pedersen

## Abstract

Cancer develops by accumulation of somatic driver mutations, which impact cellular function. Non-coding mutations in non-coding regulatory regions can now be studied genome-wide and further characterized by correlation with gene expression and clinical outcome to identify driver candidates. Using a new two-stage procedure, called ncDriver, we first screened 507 ICGC whole-genomes from ten cancer types for non-coding elements, in which mutations are both recurrent and have elevated conservation or cancer specificity. This identified 160 significant non-coding elements, including the *TERT* promoter, a well-known non-coding driver element, as well as elements associated with known cancer genes and regulatory genes (e.g., *PAX5*, *TOX3*, *PCF11*, *MAPRE3*). However, in some significant elements, mutations appear to stem from localized mutational processes rather than recurrent positive selection in some cases. To further characterize the driver potential of the identified elements and shortlist candidates, we identified elements where presence of mutations correlated significantly with expression levels (e.g. *TERT* and *CDH10*) and survival (e.g. *CDH9* and *CDH10*) in an independent set of 505 TCGA whole-genome samples. In a larger pan-cancer set of 4,128 TCGA exomes with expression profiling, we identified mutational correlation with expression for additional elements (e.g., near *GATA3*, *CDC6*, *ZNF217* and *CTCF* transcription factor binding sites). Survival analysis further pointed to *MIR122*, a known marker of poor prognosis in liver cancer. This screen for significant mutation patterns followed by correlative mutational analysis identified new individual driver candidates and suggest that some non-coding mutations recurrently affect expression and play a role in cancer development.

## Introduction

Cancer develops and progresses by accumulation of somatic mutations. However, identification and characterisation of driver mutations implicated in cancer development is challenging as they are greatly outnumbered by neutral passenger mutations^1–3^. Driver mutations increase cell proliferation, and other properties, by impacting cellular functions. Their presence is thus a result of positive selection during cancer development. Although the mutational process differs between patients, their cancer cells are subject to shared selection pressures. Driver mutations therefore recurrently hit the same cellular functions and underlying functional genomic elements across patients^4^. This allows statistical prediction of driver mutations and the cancer genes and regulatory elements they accumulate in by analysis of mutational recurrence across sets of cancer genomes^1–3^. In addition, characterization of mutations in predicted driver elements by their correlation with gene expression and patient survival can further support element cases with driver potential.

Concerted sequencing efforts and systematic statistical analysis by the International Cancer Genome Consortium (ICGC) and others have successfully catalogued protein-coding driver genes and their mutational frequency in pan-cancer and individual cancer types^5,6^. While this initial focus on protein-coding regions has dramatically expanded our knowledge of cancer genetics, the remaining 98% non-coding part of the genome has been largely unexplored. With the emergence of large sets of cancer genomes^7^, it is now possible to systematically study the role and extent of non-coding drivers in cancer development. As most non-coding functional elements are either involved in transcriptional regulation (promoters and enhancers) or post-transcriptional regulation (non-coding RNAs), non-coding drivers are expected to impact cellular function through gene regulation. A central aim of this study is therefore to systematically couple non-coding driver detection with the study of gene expression.

Few non-coding driver candidates have been identified and only a small subset has been shown to have functional or clinical consequences. The best-studied example is the *TERT* promoter, with frequent mutations in melanoma and other cancer types that increase expression in cellular assays^8,9^. A few other cases of non-coding drivers have been reported, including splice site mutations in *TP53* and *GATA3*^*10,11*^ as well as mutations in a distal *PAX5* enhancer that affect expression^12^.

Three recent studies^2,3,13^ have screened for drivers among promoters, enhancers, and individual transcription factor binding sites (TFBSs) using mutational recurrence in large sets of pan-cancer whole genomes. In combination, they report several hundred non-coding elements. The potential for affecting expression has only been studied for a subset of these. Promoter mutations were found to correlate with expression in cancer samples for *PLEKHS1*^*3*^, *SDHD*^*2*^, *BCL2*, *MYC*, *CD83*, and *WWOX*^*13*^. Melton et al. additionally identified mutations near *GP6* and between *SETD3* and *BCL11B* that reduced expression in cellular assays^2^. Negative correlation with survival was observed for promoter mutations in *SDHD*^*3*^ and *RBM5*^*13*^ for melanoma patients. Taking a different approach, Fredriksson et al. screened for expression correlation of mutations in promoters of all genes and found global significance for only *TERT*^*14*^. In addition, mutations in the *TERT* promoter were also associated with decreased survival in patients with thyroid cancer^14^.

Here, we screened for non-coding elements with surprisingly high conservation levels and cancer specificity followed by a characterization of mutation correlation with expression and survival. An extended set of regulatory element types and non-coding RNAs (ncRNAs) was created for this purpose. We developed a two-stage procedure, called ncDriver, to screen for candidate driver elements to reduce the false positive rate. In this procedure, we first identified recurrently mutated elements and then evaluated these based on combined significance of cancer type specificity and functional impact, as measured by conservation. Considering the local relative distribution of 4 mutations between positions, cancer type and conservation level, ensures robustness against mutation rate variation along the genome. Furthermore, for cancer type specificity, we estimate the expected mutation frequency given the mutation context and cancer type to account for cancer-specific mutation signatures. This approach is conceptually similar to the recent OncodriveFML method^15^. In contrast to previous studies, we included both SNVs (single nucleotide variants) and INDELs (small insertions and deletions) in the analysis. The screen identified 160 significant non-coding elements, with an enrichment of regulatory elements near known protein-coding cancer drivers. We also screened genome-wide TFBS sets for individual transcription factors to investigate whether entire transcription factor regulatory networks collectively had surprising mutational patterns and showed potential driver evidence.

To further evaluate the driver potential of significant elements, we characterized the mutations in these elements through expression perturbation using correlation of mutations in regulatory regions with gene expression levels. For this purpose we used an independent pan-cancer set of 4,128 exome capture samples with paired RNAseq samples^16^. This candidate driven approach identified significant expression correlations for individual candidates as well as for genome-wide TFBS sets, extending observations by Fredriksson et al.^14^. We further evaluated the association of mutations in significant elements with patient survival. Though limited by small numbers of patients mutated for individual elements, this analysis identified candidate drivers and mutations of potential clinical relevance, including liver cancer mutations of the poor prognosis biomarker microRNA (miRNA) *MIR122*.

## Results

### Pan-cancer screen for non-coding elements with conserved and cancer specific mutations

To screen for non-coding elements with elevated conservation and cancer specificity, we used a set of 3.4M SNVs and 214K INDELs from a previous study of 507 whole-cancer-genomes from ten different cancer types (**Supplementary Table 1**)^7^. Mutation rates varied more than five orders of magnitude across samples, with the number of SNVs per sample (median=1,988) about nine times higher than for INDELs (median=198; Fig. 1a). More than ten million non-coding elements spanning 26% of the genome collected from ENCODE and GENCODE were screened, including long ncRNAs (lncRNAs), short ncRNAs (sncRNAs), pseudogenes, promoters, DNaseI Hypersensitive Sites (DHSs), enhancers and TFBSs (Fig. 1b,c)^17,18^. Protein-coding genes (n=20,020; 1.1 % span) were included as a positive control.

**Figure 1.**
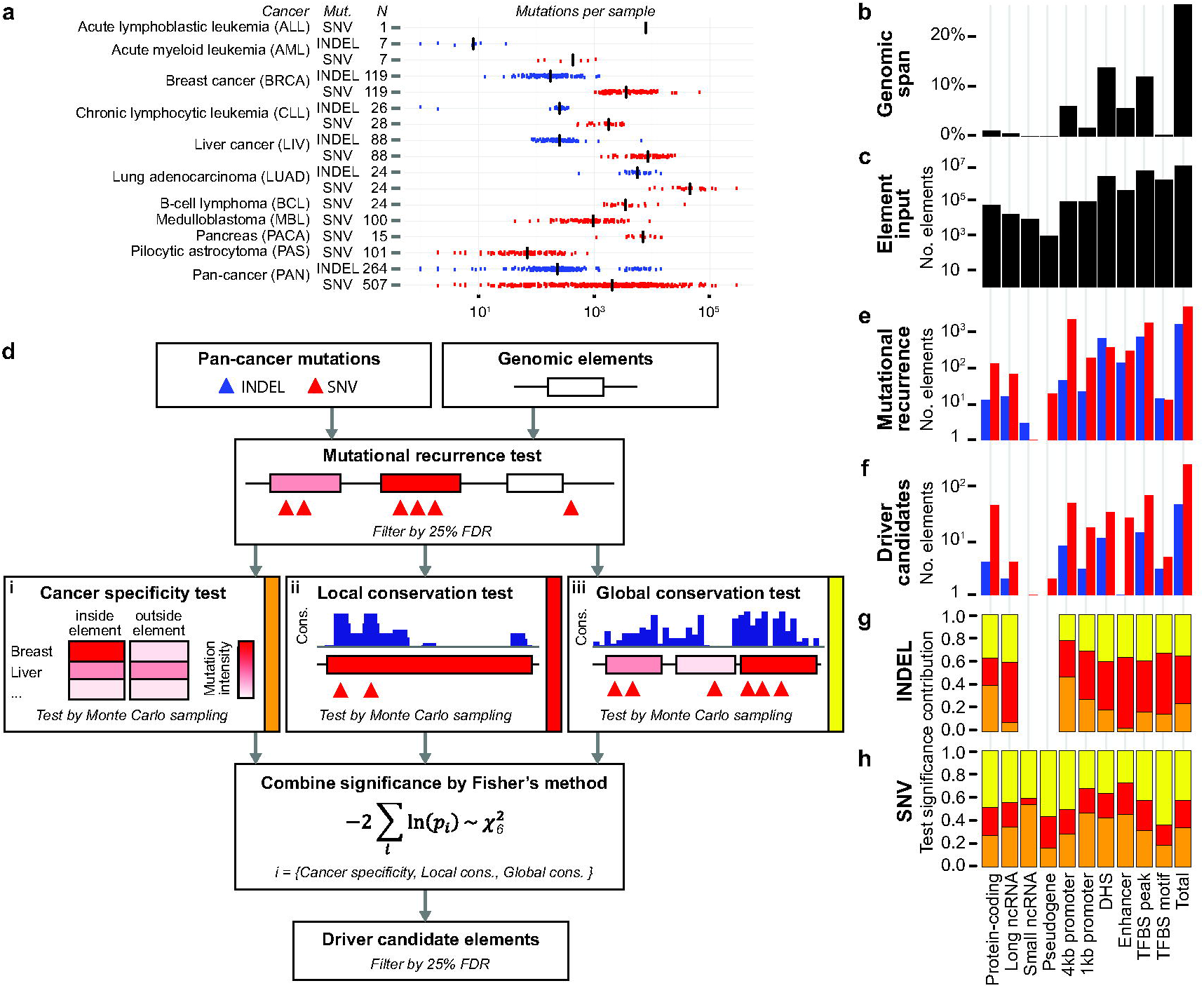
Overview of the two-stage procedure detecting for non-coding elements with cancer-specific and conserved mutations and its application to a pan-cancer whole-genome data set. (a) Summary of the input data, showing the cancer type (Cancer), mutation type (Mut.), number of samples (N) and number of mutations per sample in the whole-genome data set^7^. SNVs are indicated by red color, INDELs by blue color and the median number of mutations is indicated with a black bar. (b-c) Genomic span and count of input elements for each element type. (d) Workflow of ncDriver, a two-stage procedure for non-coding driver detection. Elements passing the *Mutational recurrence test* of the first stage are passed on to the second stage tests *Cancer specificity test* (i), *Local conservation test* (ii) and *Global conservation test* (iii). (e) Counts of elements that passed the *Mutational recurrence test* at a 25% FDR threshold for SNVs (red) and INDELs (blue). (f) Counts of significant elements that passed the combined significance using Fisher’s method and 25% FDR threshold. (g-h) Relative contribution of the *Cancer specificity test* (i; orange), *Local conservation test* (ii; red), *Global conservation test* (iii: yellow) to the combined significance of the significant elements of each element type for INDELs and SNVs.

Each element type was separately screened using a new two-stage procedure, called ncDriver (Fig. 1d). Its underlying idea is to restrict the element selection (second stage) to tests that are robust to the variation in the mutation rate^1^ and thereby reduce the false positive rate. These tests evaluate the relative distribution of mutations instead of the overall number of mutations. More specifically, these tests consider the cancer-type specific mutational processes and sequence context preferences, when evaluating cancer specificity, and evaluate mutations enriched for conserved and functional sites. This is conceptually similar to test of positive selection for protein-coding regions that evaluate the enrichment of amino-acid changing substitutions over silent ones^19^. To reduce the number of tests performed and focus on relevant elements with enough mutations for the tests to be powerful, we first identified elements with mutational recurrence (first stage) and among these we evaluate the actual driver significance using a combination of cancer specificity and conservation (second stage).

In more detail, first, a lenient test of mutational recurrence identified a total of 6,529 elements (n_SNV_=4,908, n_INDEL_=1,621) with elevated mutation rates (Fig. 1e). Second, for each element type the recurrently mutated elements were passed on to three separate driver tests for candidate selection. Each of these tests address different aspects of the mutations’ distribution. *Cancer specificity test:* Based on previous observations of cancer specificity of known protein-coding drivers^5^, we evaluated if the mutations within each element showed a surprising cancer specific distribution given the cancer specific mutational signatures (Fig. 1d.i). *Local conservation test:* Since it is often not understood how function is encoded in non-coding elements, we used evolutionary conservation as a generic measure of functional importance. We tested if mutations showed a surprising preference for highly conserved positions within each element, which suggests that mutations of functional impact are enriched and have been selected for (Fig. 1d.ii). *Global conservation test:* As highly conserved elements are more likely to be key regulators^18^, we also tested if the conservation level of mutated positions in a given element was surprisingly high compared to the overall conservation distribution across all elements of the same type (Fig. 1d.iii). Finally, we used Fisher’s method to combine the significance of the cancer specificity and conservation tests and q-values (q) were used to threshold (25% false discovery rate; FDR) and rank the final lists for each element type for a total of 295 significant elements (Fig. 1f; **Supplementary Table 2**). The final selection is thus based on a combination of three different aspects of the mutations distribution, given the cancer type specific mutational signatures, to improve overall driver detection power.

For the final set, the most significant element was preferred when overlap occurred, which resulted in 160 unique non-coding elements and 48 protein-coding genes. Of these, 35% (39 of 208) were found based on INDELs, despite they only comprise 4% of the full mutation set (Fig. 1f). The contribution of the three different driver tests to the significance of the final candidates varied among element and mutation types (Fig. 1g,h). Generally, the Local conservation test made the largest contribution for INDELs and the Global conservation test made the largest contribution for SNVs. The contribution of the cancer specificity test was largest for sncRNAs called by SNVs.

For protein-coding genes, known cancer-drivers in COSMIC^6^ are top-ranked and enriched among significant elements for both the SNV set (13.0x; p-value=p=2.4x10^−9^) and the INDEL set (102.6x; p=9.1x10^−5^; **Supplementary Table 3**)^6^. If applied individually, all three driver tests also resulted in enrichment of known protein-coding drivers, with 34.6x enrichment for the cancer specificity test (p=4.8x10^−11^), 17.1x for the local conservation test (p=1.7x10^−3^), and 10.6x for the global conservation test (p=6.5x10^−8^; **Supplementary Table 3**). All three tests are thus able to detect signals from known protein-coding drivers, despite not tailored for this purpose.

To further evaluate driver evidence for both individually identified elements and the set as a whole, we asked if an independent data set supported the findings. For this, we applied ncDriver specifically to the above defined set of 208 significant elements using another set of 505 whole-genomes from 14 cancer types^14^ (**Supplementary Fig. 1**). Even for true drivers, we only expected limited recall of individual non-coding elements as the two sets differ in their cancer-type composition affecting the statistical power to recall cancer-type specific drivers. Furthermore, the available whole genome data sets generally have limited statistical power to detect true drivers with only few driver mutations and hence small effect sizes. Such drivers are unlikely to be consistently detected across sets, known as winner’s curse^20^.

Overall 17 elements were recalled (**Supplementary Table 2**), including eight protein-coding genes (*TP53*, *KRAS*, *FBXW7*, *PIK3CA*, *TMEM132C*, *CSMD1*, *BRINP3* and *CDH10*), one enhancer (associated with the known *TERT* promoter sites^8,9^), two protein-coding gene promoters (*CDH10* and *MEF2C)*, three lncRNA promoters (*RP11-760D2.11*, *RP11-805F19.1*, and *RP11-463J17.1*), two TFBS peaks (associated with *PFKP* and *MROH1*) and one TFBS motif associated with FSHR (**Supplementary Fig. 3**). The overall number of elements recalled is six times higher than expected by chance (**Supplementary Table 4**; p=0.001; Monte Carlo test, see Methods). Among the element types, where any number of elements were recalled, we identified three element types with significant enrichment (p<0.003) (**Supplementary Table 4**).

A given driver gene may be affected by mutations at different nearby regulatory elements. We therefore performed another recall analysis, using the same independent dataset, in which we extended the element set to include all elements associated with the same genes as our elements (n=208). We analyzed this extended set using the original approach to screen for possible driver evidence in the independent set of cancer genomes (**Supplementary Fig. 1**). For this we screened 251,333 elements (2.3% of all input elements) associated with these 208 genes. At the *gene level*, 82 genes were recalled by one or more non-coding elements, with only three called by evidence in the protein-coding gene itself (Fig. 2a; **Supplementary table 2**). The recall rate was a bit higher for known cancer genes^6^ (48%; 11 of 23) than for other genes (37%; 68 of 185), though not significant (p=0.36; Fishers’ exact test).

**Figure 2.**
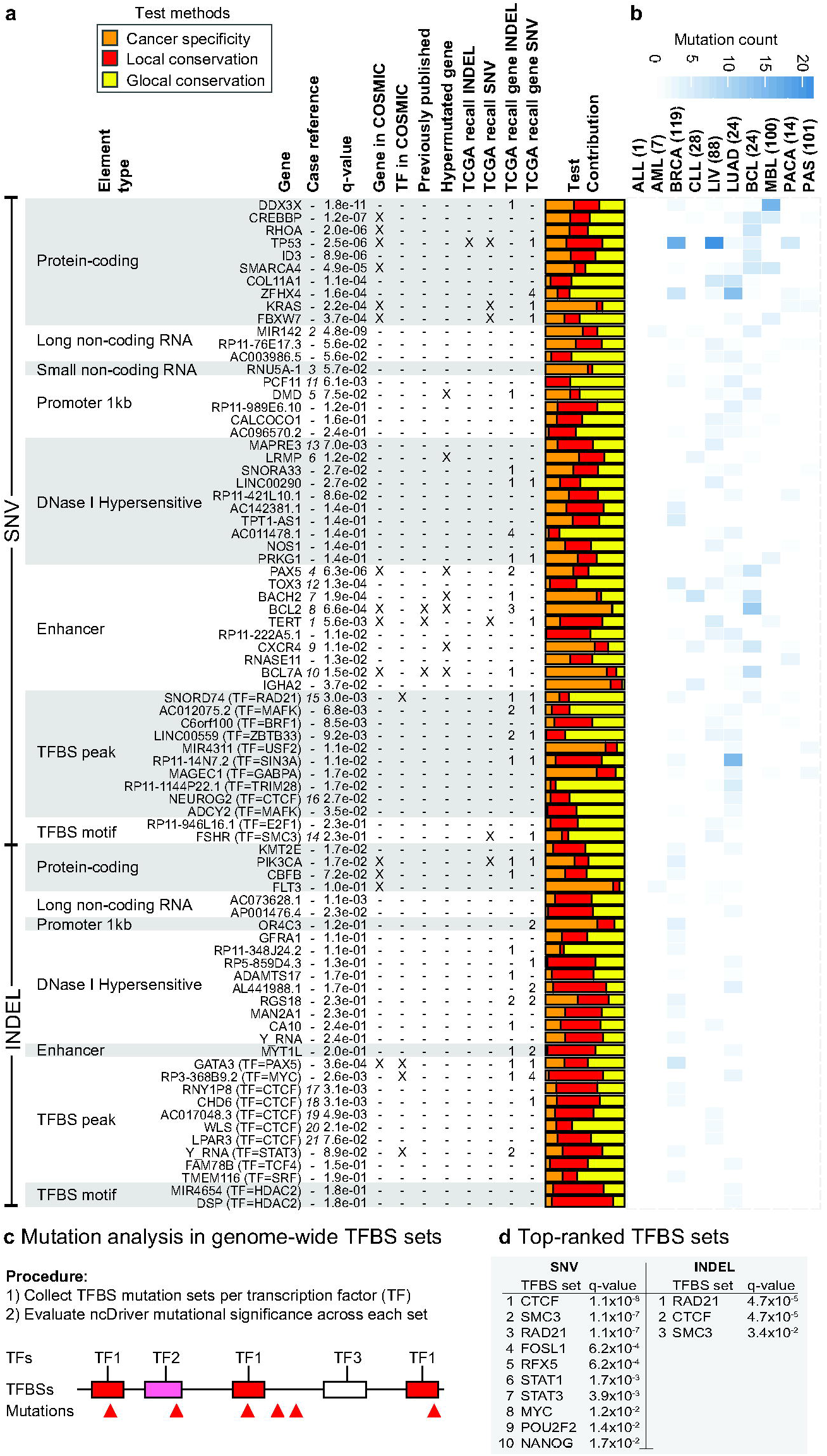
Top-ranked significant non-coding elements from pan-cancer driver screen. (a) Table with top-ten significant elements for each element type for both SNVs and INDELs ranked by combined significance. *Gene*: Gene name or name of gene with nearest transcription start site in case of regulatory elements (DHS, enhancers, and TFBS). *Case reference:* Reference number of specific cases. *q-value:* ncDriver combined significance using Fisher’s method and Benjamini-Hochberg corrected for each element type. *Gene in COSMIC:* Gene name present in COSMIC database of known drivers^6^. *TF in COSMIC:* Transcription factor of TFBS element present in COSMIC. *Previously published*: Element is overlapping a region found in previously published non-coding driver screens^2,3^. Only most significant element retained when elements overlap between element types. *Hypermutated gene*: Gene name previously characterized as a hypermutated gene^22^. *TCGA recall INDEL/SNV:* Individual element recalled in TCGA independent whole-genome data set^14^. *TCGA recall gene INDEL/SNV:* Number of elements recalled at the gene level in TCGA independent whole-genome data set. *Test Contribution:* Relative contribution of Cancer specificity test (orange), Local conservation test (red) and Global conservation test (yellow) to the combined significance using Fisher’s method. (b) Heatmap of mutation count per cancer type. Cancer type abbreviations defined in Fig. 1. Pseudogenes and 4 kb promoters are listed in Supplementary Table 2. (c) Overview of the procedure for mutation significance analysis in TFBS sets for individual transcription factors. (d) The top-ranked significant TFBS sets, denoted by their transcription factor, for SNVs and INDELs.

We were able to recall known cancer drivers in the independent data set of cancer genomes. However, the relatively low number of recalled elements (17 out of 208) indicates that there are few non-coding drivers with high pan-cancer mutations rates and potentially a presence of false positives.

### Significant non-coding elements identified in the pan-cancer screen

Significant non-coding elements were found in all element types, though in varying number and significance, with most for TFBS peaks (TFP; n_SNV_=68; n_INDEL_=14) and least for sncRNAs (nSNV=1) (Fig. 2a; **Supplementary Table 2**). The non-coding regulatory elements are annotated by their nearest protein-coding gene. Overall, the significant non-coding (regulatory) elements show an enriched (4.6x) association with known cancer driver genes (14 of 121; p=8.6x10^−6^; **Supplementary Table 3**). The highest enrichments are seen for promoters (14.7x; p=1.5x10^−5^) and enhancers (16.2x; p=2.9x10^−7^).

The significant elements include the well-studied *TERT* promoter region (**Supplementary Table 2**)^8,9^. As an overlapping enhancer element achieves higher significance, it represents the region in the final list (Fig. 2a.*1*, i.e., case *1* in column three in Figure 2a). Several candidates from previous screens are also present (n=5; **Supplementary Table 2)**^2,3^.

The primary miRNA transcript *MIR142*, a lncRNA, is the most significant non-coding driver candidate overall (q=4.8x10^−9^; **Fig 2a.*2*; Supplementary Fig. 2a,b**). Ten SNVs from AML, CLL, and BCL lymphomas fall in the 1.6kb-long transcript. Three of these hit the highly conserved precursor miRNA region (88 bp), which forms a hairpin RNA structure, potentially directly affecting the biogenesis of the mature miRNA. While SNVs in the miRNA precursor were previously reported for AML and CLL^12,21^, we here find SNVs across the entire primary miRNA and for all three haematological types (Fig. 2b). Apart from an uncharacterized lncRNA (*RP11-76E17*), a U5 spliceosomal RNA (*RNU5A-1*; **Fig 2a.*3*; Supplementary Fig. 2c,d**), and two pseudogenes (**Supplementary Table 2**), the remaining non-coding elements are gene regulatory. A distant enhancer of the B-cell specific transcription factor *PAX5* was recently found to be recurrently mutated in CLL and other leukemias with an effect on expression^12^. Here we detect an overlapping TFBS peak for *RAD21*, associated with the non-coding gene *RP11-397D12.4*, with four SNVs in both of CLL and BCL (q=7.2x10^−2^; Fig. 3a,b). In addition, our top-ranked enhancer element is located within the first intron of *PAX5* and hit by eight SNVs in BCL and two in LUAD (q=6.3x10^−6^; Fig. 2a.*4*; Fig. 3c). Interestingly, five of the mutations fall within a TFBS for CTCF (q=2.4x10^−4^; Fig. 3c).

**Figure 3.**
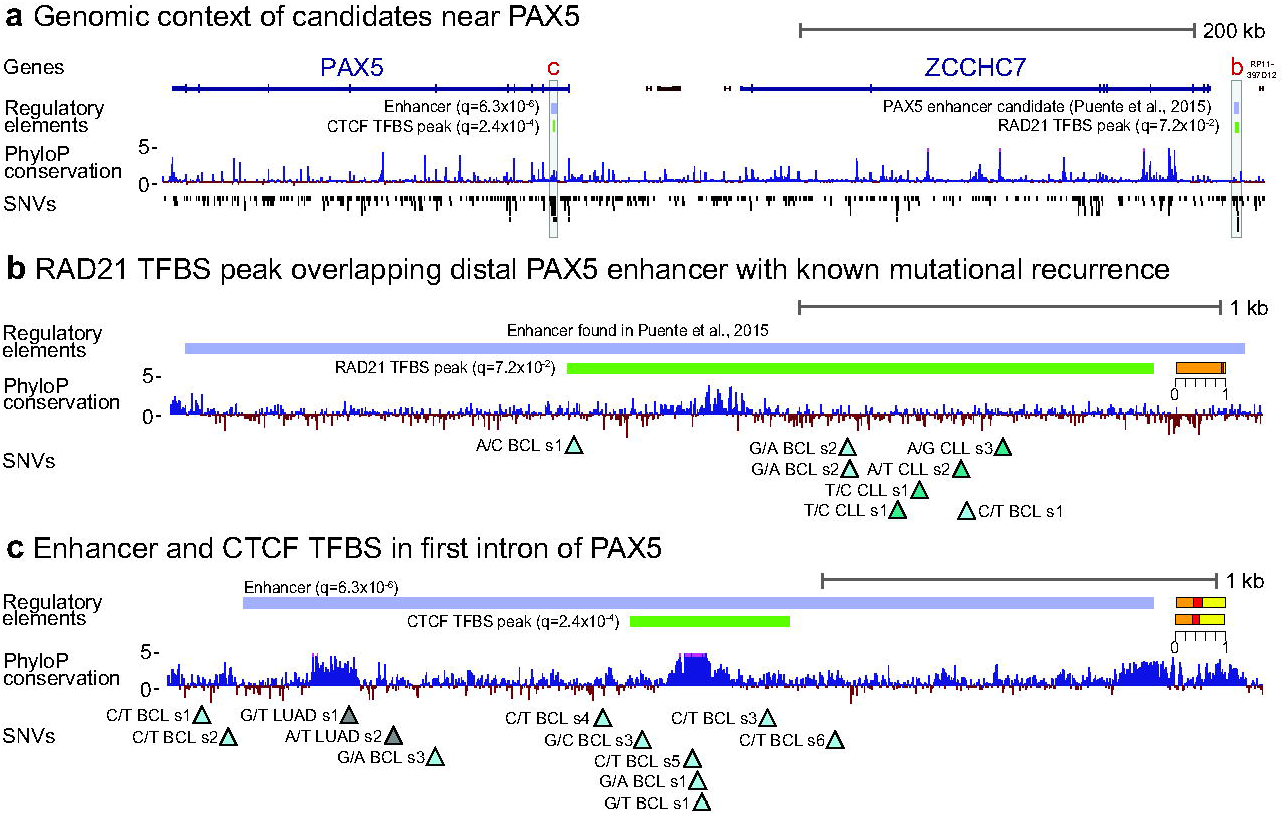
Significant regulatory elements associated with *PAX5*. (a) Genomic context of *PAX5* with protein-coding genes (blue), non-coding genes (brown), significant regulatory elements, PhyloP conservation and SNVs. (b) The element *RAD21* TFBS peak (Supplementary Table 2) overlaps an enhancer with known mutational recurrence and effect on *PAX5* expression^12^. Mutations (triangles) are annotated with nucleotide change (from/to), cancer type (abbreviation and color), and sample number (s1-k). The relative significance-contribution from each of the three mutational distribution tests shown as in Fig 2a. (The same applies to the other case illustrations.) (c) Regulatory elements in the first intron of *PAX5*. Both enhancer and *CTCF* peaks are individually significant with contributions from the conservation tests.

Among the SNV-top-ranked promoters (*DMD*), DHS elements (*LRMP*) and enhancers (*PAX5, BACH2*, *BCL2*, *CXCR4*, and *BCL7A*) are highly cancer type specific cases with many BCL or CLL mutations (Fig. 2a.*4-10,b*; Fig. 3). These are known targets of somatic hypermutations affected either through translocations to Immunoglobulin loci (e.g., *BCL2* and *PAX5*) or by aberrant somatic hypermutations targeting transcription start site regions of genes highly expressed in the germinal centre (e.g., *DMD* and *CRCX4*)^12,22,23^. However, the conservation tests show a non-random mutation pattern for some of these (*PAX5* and *DMD* in particular), suggesting an effect of selection and driver mutations.

Among promoters, the 3’-end processing and transcription termination factor *PCF11* is ranked first by SNVs. It is is hit by seven SNVs (q=6.2x10^−3^) from breast, lung and liver cancer types (**Supplementary Table 2**) in its 5’UTR, which has a high density of transcription factor binding sites^18,24^. The mutations are biased toward highly conserved positions, as evidenced by the conservation test contributions (Fig. 2a.*11*; Fig. 4a). Downregulation of *PCF11* affects both 25 26,27 transcription termination^25^ as well as the rate of transcription re-initiation at gene loops^26,27^. Mutational perturbation of *PCF11* may thereby affect transcriptional regulation.

**Figure 4.**
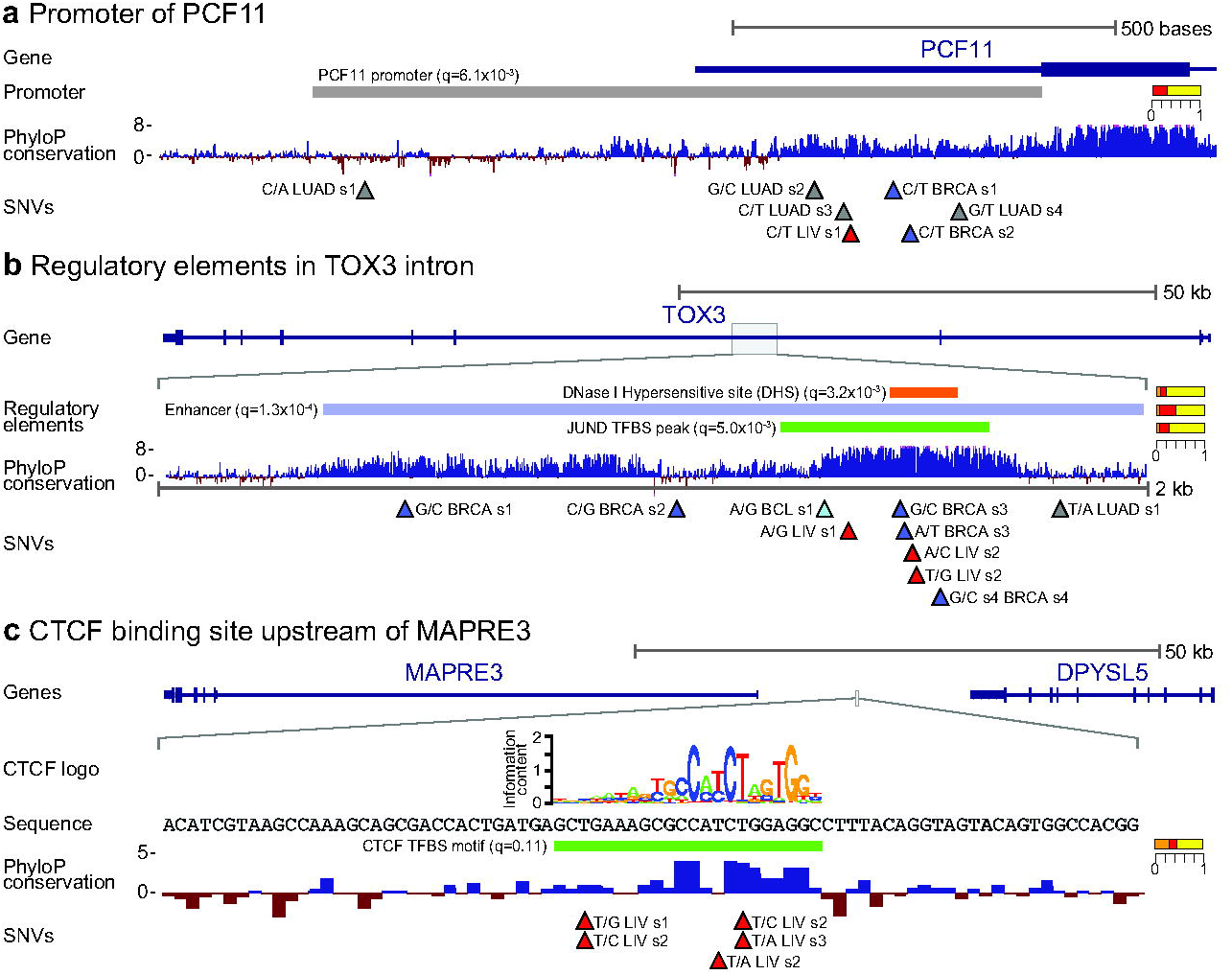
Cases of significant regulatory elements. Top rows show the genomic context with nearby gene and rows below show detailed views of the regulatory elements, PhyloP conservation scores, and SNVs. SNV annotations and color-scheme as in Fig. 3. (a) Mutations in the significant upstream promoter element of *PCF11*. (b) Mutations in significant intronic elements of *TOX3*. The three elements achieve similar combined significance after multiple testing correction. (c) Mutations in the significant *CTCF* TFBS element upstream of *MAPRE3*. The *CTCF* sequence logo and nucleotide sequence of the region is shown.

A 1.9Kb-long enhancer in an intron of *TOX3* is ranked second by SNVs and also achieves significance primarily from the conservation tests (Fig. 2a.*12*; Fig. 4b). It is hit by ten SNVs (q=1.3x10^−4^) in breast, liver, lung, and BCL cancer types. Numerous TFBS peaks overlap the mutations, with a *JUND* TFBS achieving the highest individual significance (q=5.0x10^−3^). *TOX3* is involved in bending and unwinding of DNA and alteration of chromatin structure^28^. It is a known risk gene for breast cancer^29^, where it is also somatically mutated at a moderate rate^30^. In line with this, we observed the most SNVs in breast cancer (n=5).

The SNV-top-ranked DHS element (q=7.0x10^−3^) is located upstream of the *MAPRE3* gene (Fig. 2a.*13*; Fig. 4c). It is hit by five mutations in liver cancer, which also overlap a TFBS for *CTCF*(q=0.1). The lower final significance of the TFBS than the DHS elements is a result of the multiple testing correction procedure. There is high mutational recurrence for the *CTCF* TFBS (q=1.9x10^−3^). The *MAPRE3* gene is microtubule associated, with frameshift mutations reported for gastric and colorectal cancers^31^.

The SNV-top-ranked SMC3 TFBS motif downstream of FSHR provides a similar example of a previously unknown recurrently mutated TFBS with three liver cancer mutations and three additional SNVs located just outside the element (Fig 2a.14; **Supplementary Fig. 3**).

Overall a large fraction of the candidate TFBSs from both SNVs and INDELs are either *CTCF*, *RAD21*, or *SMC3* binding sites (25 of 91; **Supplementary Table 2**; Fig. 2a.*14-21*), which are all 32 part of the cohesin complex^32^. Recently, an elevated SNV rate at binding sites of the cohesin complex have been reported for several cancer types by others^33,34^. The cohesin complex is a key player in formation and maintenance of topological chromatin domains^35,36^, suggesting that non coding mutations could play a role shaping the chromatin structure during cancer development. On the other hand, the elevated mutation rate could be caused by the specific environment induced by the binding of transcription factors^37^.

The large fraction of significant cohesin binding sites suggests that binding sites of some transcription factors (TFs) may be overall more mutated and perturbed than others in cancer development. To answer this, we screened genome-wide sets of TF binding site motifs (ntotal=1.7M) for individual TFs (109 TFs comprising 915 individual subtypes) from ENCODE that are found within a TF peak^38^ for overall driver evidence using the ncDriver approach. As the number of hypotheses is smaller than for the above screen of individual elements, we did not apply the initial mutation recurrence filter (**Supplementary Note 1**).

This identified transcription factors with significant binding site sets for both SNVs (n=25) and INDELS (n=4; q<0.05; Fig. 2d; **Supplementary Table 5**). The genes associated with the mutated sites are enriched for functional terms related to cancer for seven of the top-ranked TFBS sets (**Supplementary Table 6**). The cohesin complex members (*CTCF*, *RAD21*, and *SMC3*) were top-ranked for both SNVs (q<1.1x10^−7^) and INDELs (q<3.4x10^−2^; Fig. 2d). We further performed a genome-wide analysis of the mutations in *CTCF* binding sites to investigate their functional properties, focussing on the binding sites of the most common subtype (subtype descriptor 1; disc1) (**Supplementary Note 2**). Together, our results show that the mutation rate is elevated at highly conserved and high affinity *CTCF* binding sites in active, open-chromatin regions^39^ (**Supplementary Fig. 4**). The increase in mutation rate not only at functionally important sites (position 16), but also at apparently non-functional sites (3’ flanking region), suggests that much of the increase may be driven by mutational mechanisms caused by micro-environmental conditions coupled to *CTCF* binding. Specifically, spacer DNA regions between the core *CTCF* binding site and flanking optional binding sites appear to be physically bent during binding^40,41^, which may affect mutation rates.

### Correlation of mutations in significant non-coding elements with gene expression

Mutations in non-coding elements may affect gene expression and thereby cellular function, exemplified by mutations in the *TERT* promoter^8,9,14^. The effect may be caused by various mechanisms, including perturbation of transcription initiation^8,9^, chromatin structure^42^, and post-transcriptional regulation^43^. The potential for mutations in elements impacting cellular function can be evaluated by analyzing differences in gene expression. We therefore developed a pan-cancer test for mutations correlating with increased or decreased gene expression levels and applied it to a large independent expression dataset from TCGA (Fig. 5a-f). Though we cannot evaluate whether the mutations cause expression difference, significant expression correlation can help identify and prioritize driver candidates and lead to specific functional hypotheses.

**Figure 5.**
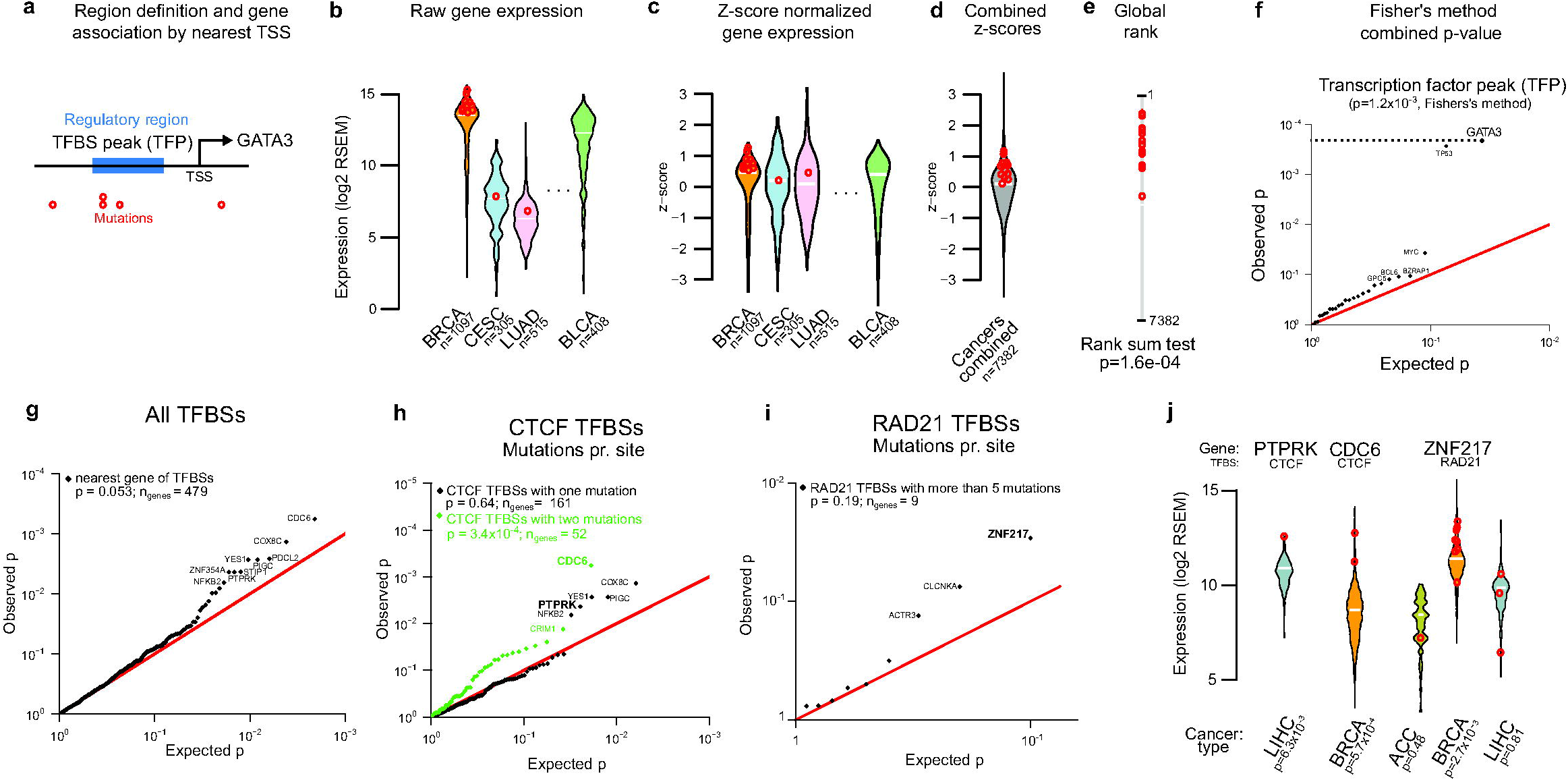
Test method and correlation analysis of mutations in significant non-coding elements with gene expression. (a-f) Overview of expression correlation test, exemplified by *GATA3* and the set of significant TFBS peak elements (TFPs). (a) Elements are associated to genes using the nearest TSS. (b) Raw expression levels (log2 RSEM) are obtained for 7,382 samples across 22 cancer types and mutated samples are identified. (c) Expression levels are z-score normalized within each cancer type and (d) combined. (e) The p-value of the mutated samples in the distribution of the combined z-score-ranked set is found using a rank-sum test. (f) P-values of significant elements and their associated genes are shown in a qq-plot with *GATA3* highlighted. The red line indicates expected p-values under the null hypothesis of no expression correlation. The combined p-value of the correlation between mutations and expression levels across the set of candidate regions is found using Fisher’s method. Cancer type abbreviations: Lung adenocarcinoma (LUAD), breast cancer (BRCA), bladder cancer (BLCA), cervical squamous cell carcinoma (CESC). (g) Gene-expression correlation for all mutations (both SNVs and INDELs) in significant TFBS sets. Rank-sum test p-values of individual genes are shown as qq-plot. Combined significance across all genes is found using Fisher’s method and shown in upper left corner (similarly for panels h and i). (h) Expression correlation for *CTCF* TFBSs mutated once (black) or twice (green). The combination of p-values was done separately for the set of TFBSs mutated once and twice. (i) Expression correlation for *RAD21* TFBSs mutated more than five times. (j) Examples of mutated TFBSs and their associated gene-expression distributions in individual cancer types (exemplified genes emphasized in (h) and (i)). Expression levels of mutated samples are shown (red circles). The expression correlation significance within each individual cancers type is given below the plot. Cancer type abbreviations: Liver hepatocarcinoma (LIHC), breast cancer (BRCA), Adrenocortical carcinoma (ACC).

**Figure 6.**
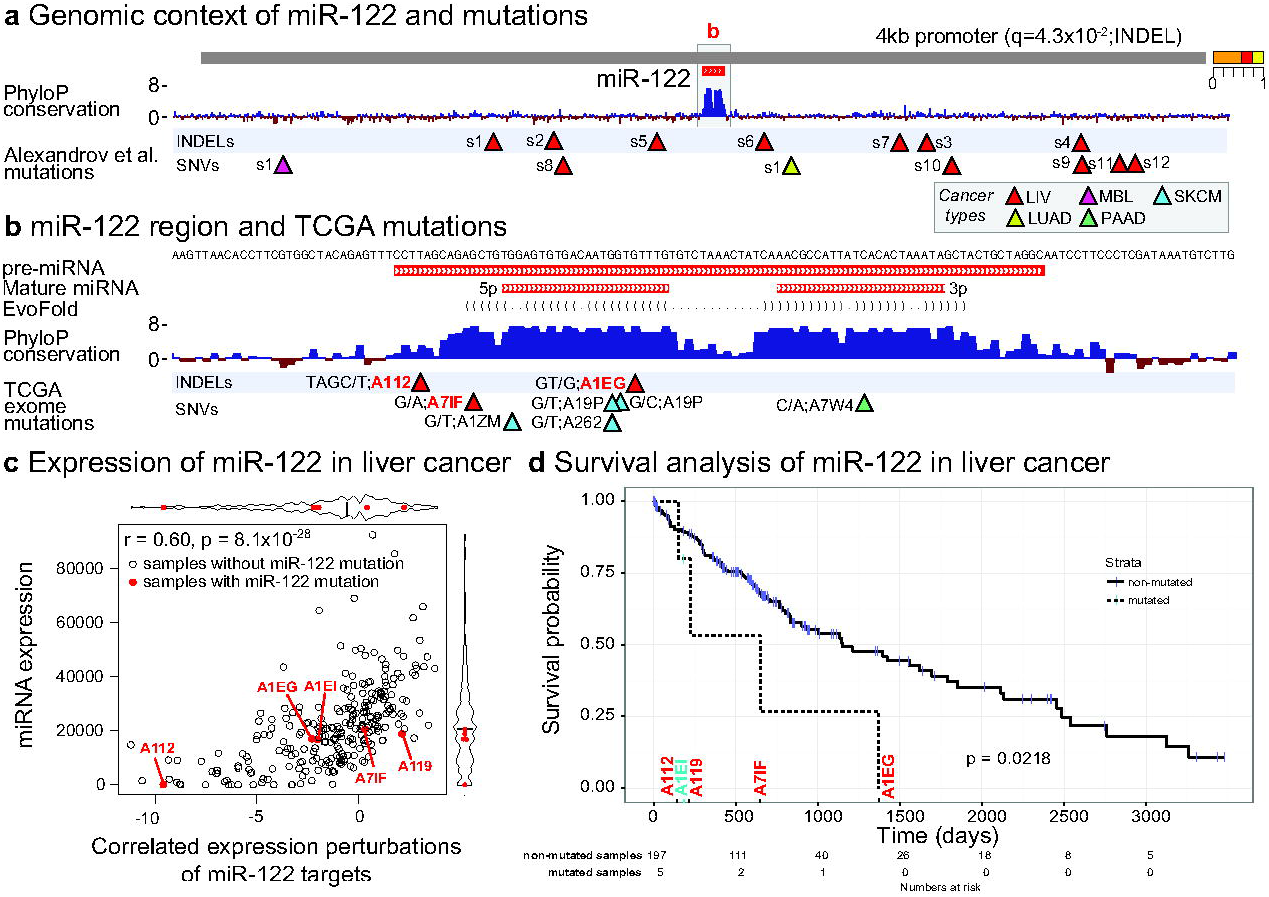
Mutations in driver candidate miR-122 and their correlation with expression and survival. (a) The 4kb genomic region of the *MIR122* gene detected as a significant element in the driver screen of the original data set with PhyloP conservation scores, INDELs and SNVs. Cancer types are color coded in the grey shaded box. (b) Close up of the miR-122 region with tracks for pre-miRNA, mature miRNA, EvoFold secondary structure prediction, PhyloP conservation scores and exome mutations from TCGA. Mutations are named by their associated sample ID and colored red if used later in the correlation analysis of expression and survival shown in c and d. (c) Correlation between miR-122 expression and miR-122 target site motif enrichment in 266 TCGA liver cancer samples. Motif enrichment is based on expression of mRNAs and motif occurrences in their 3’UTRs (see Methods). Samples mutated in the miR-122 region in (b) are indicated in red. (d) Survival correlation analysis of TCGA liver mutations in miR-122. The number of mutated samples and non-mutated samples at each time point is indicated below the plot.

Essentially, the idea is to first make expression levels comparable across cancer types by applying z-score normalization to the expression values for a given gene within each cancer type **(Fig. 5b,c)**. Then evaluate differences between mutated samples and non-mutated samples combined across cancer types, using a non-parametric rank sum test **(Fig. 5d,e)**. Finally, where relevant, combine such statistical evidence across all the genes regulated by a given set of non-coding elements, e.g. all TFP elements found significant in the driver analysis **(Fig. 5f)**. Each tested element was associated to the nearest gene, and the test was based on gene expression in an independent set of 7,382 RNAseq samples of which 4,128 had paired exome mutation calls (both SNVs and INDELs)^16^. Though the power to call mutations from exome capture data is highest in protein-coding regions, 50% of the calls are found in the non-coding part of the genome.

We first focused on sets of elements with regulatory potential, and thus evaluated correlation effects in TFBS peak, 1kb promoter and DHS element types. Mutations in the set of TFBS peak candidates correlated overall with unusual expression levels (p=1.2x10^−3^; Fig. 5f). The significant expression correlation was primarily driven by mutations at two known cancer drivers TP53 (p=2.3x10^−4^) and GATA3 (p=2.1x10^−4^), with MYC also nominally significant (p=3.7x10^−2^). The promoter and DHS candidate sets did not achieve overall significance (**Supplementary Fig. 6**). The GATA3 mutations (n=15) all reside in intron four of the gene and most are INDELs from breast cancer (n=11) that disrupt the acceptor splice site, which leads to abnormal splicing and codon frame shift as described previously for the luminal-A subtype of breast cancer^10,11^. In addition, one lung adenoma SNV also disrupt the splice site. The association between *GATA3* splice-site mutations and higher *GATA3* expression is, to our knowledge, novel. Similarly, most of the *TP53* mutations affect splice-sites in intron eight. Both germline and somatic driver mutations in splice sites are known for *TP53*^*44,45*^.The *GATA3* and *TP53* results show that the expression test can identify known non-coding driver mutations that correlate with transcript abundance.

We next focused on the effect of TFBS mutations on nearby gene expression. For this, we applied the expression test to the 29 significant TFBS sets (**Supplementary Table 5**) and subsets thereof as indicated in Fig 5n,i. In combination, the expression correlation of the full set of TFBS mutations showed borderline significance (p=0.053; Fig. 5g), with a limited set of genes that deviate from the expected p-values.

Both passenger and driver mutations may impact expression. As it is unlikely that passenger mutations hit the same TFBS twice by chance, we expect enrichment for true drivers among those that do. To further pursue this idea and enrich for driver mutations, we analyzed expression correlation separately for different numbers of pan-cancer mutations hitting the same type of TFBS. For most TFBS sets, the stratified subsets became small and we therefore focused on the large *CTCF* set (Fig. 5h). Overall, the set of double-hit mutations had a much stronger correlation with expression (p=3.4x10^−4^) than single-hit mutations (p=0.64). For double-hit mutations, the majority shows a deviation from the expectation, whereas for single-hit mutations, this is only the case for the five most significant genes (Fig. 5h). This shows a generally stronger correlation and a larger potential for cellular impact for double-hit than single-hit mutations, consistent with an enrichment of true drivers. To rule out that the difference was simply an effect of additional power for the double-hit mutations, we confirmed that p-values for individual double-hit mutations were generally smaller than single hit mutations (p=0.01; one sided rank sum test).

Among the individual genes top-ranked by the expression correlation analysis are well-studied cancer genes, often with tissue-specific mutation-patterns. *CDC6*, which is found in the COSMIC Gene Census database^6^, is top-ranked for all TFBS’s and also for the *CTCF* double-hit mutations (Fig. 5g,h), with two mutations in breast cancer (Fig. 5j). *CDC6* is a necessary component of the pre-replication complex at origins of replication and involved in cell-cycle progression-control via a mitotic checkpoint^46^. It mediates oncogenic activity through repression of the *INK4*/*ARF* tumor suppressor pathway^47^ and is an activator of oncogenic senescence^48^. In breast cancer, its expression correlates with poor prognosis^49^. *PTPRK* is among the few *CTCF* TFBS single-hit genes with unexpected expression correlation, with a single mutation in liver cancer (Fig. 5h,j). It is a tyrosine phosphatase associated with several cancer types^50,51^. Four liver cancer mutations in an associated *YY1* TFBS of *PTPRK* also correlate positively with expression (p=2.7x10^−2^). Individual TFBSs are hit by more than five mutations in numerous cases (n=154). Though recurrent technical artifacts may underlie most of these extreme cases, some exhibit convincing expression correlations (Fig. 5i). One such example is *ZNF217*, which is hit in an associated *RAD21* binding site by eight breast cancer mutations and by four in other cancer types. The breast cancer mutations correlate strongly with increased expression level (p=2.7x10^−3^; Fig. 5j). *ZNF217* is well studied in cancer^52^. It is a known breast cancer oncogene and an expression marker for poor prognosis and metastases development^53^. Given this, it would be a natural candidate for further studies of the clinical relevance of regulatory mutations once larger data sets become available.

### Association of mutations in significant non-coding elements with patient survival

Driver mutations may affect not only cancer development, but also cell proliferation, immune evasion, metastatic potential, therapy resistance, etc, and thereby disease progression and potentially clinical outcome^54^. An association between candidate driver mutations and clinical outcome would therefore support a functional impact on cancer biology as well as point to a potential as clinical biomarker.

To pursue this, we focused on the TCGA whole genome and exome data sets where we have information on patient overall survival time (**Supplementary Tables 10 and 11**). For the exome data set, we evaluated all candidate elements found in the original driver screen (n=208), whereas we restricted the focus to the subset of recalled elements (n=17) for the smaller, less well-powered whole genomes data set (**Supplementary Fig. 1**). For each candidate element, we restricted the focus to cancer types with at least three mutations, to retain statistical power. For each cancer type, we asked whether the patients with a mutation in the element had significantly decreased overall survival compared to patients without a mutation using a one-sided log-rank test on the Kaplan-Meier estimator. We tested the one-sided hypothesis, as we were interested in driver mutations and hence cases with worsening survival, adopting a previously discussed test strategy^55^. For an overall pan-cancer measure of significance, we combined the p-values of the individual cancer types, using Fisher’s method. Finally, elements with an estimated FDR of less than 25% were considered significant, which resulted in three protein-coding genes across both data sets and four non-coding elements based on exomes only (**Supplementary Tables 12-15**).

For protein-coding genes, *TP53* and *KRAS* were independently found to be significant in both the exome and whole-genome data sets (**Supplementary Tables 12 and 14**), with nominal significance (p<0.05) in a range of individual cancer types (**Supplementary Fig. 7 a,b, f:j**) in line with the literature ^56,57^. In addition, *NRXN1* was found significant in the exome set (q=0.09), with nominal significance (p<0.02) for the breast cancer, liver hepatocellular carcinoma, and thyroid cancer types (**Supplementary Fig. 7 c,d,e**). Though *NRXN1* has not previously been described as a driver, it is a known recurrent target of hepatitis B virus DNA integration in liver hepatocellular carcinoma^58^.

For non-coding elements, enhancer nearby *TERT* is ranked first in the whole genome data set with near-significance (q=0.32; **Supplementary Tables 15**). The highest significance for individual cancer types is seen for glioblastomas (p=0.057) and thyroid cancer (p=0.063), which are also the cancer types where *TERT* promoter mutations have previously been shown to correlate with cancer progression^59,60^.

The top-ranked non-coding element is a promoter of lncRNA *LINC00879* (q=1.6x10^−6^), with nominal significance in esophageal cancer (p=0.013) and liver hepatocellular carcinoma (p=1.5x10^−10^) (**Supplementary Fig. 8a,b**). The lncRNA is uncharacterized. Its promoter region overlaps the pseudogene *WDR82P1*. The promoter of the kinase *SGK1* is second-ranked (q=0.22), with nominal significance in stomach cancer (p=0.0002; **Supplementary Fig. 8f**). *SGK1* is overexpressed in epithelial tumours and recently associated with resistance to chemotherapy and radiotherapy^61^.

A transcription factor peak near *PCDH10* is ranked fourth (q=0.22; **Supplementary Fig. 8c**). *PCDH10* is a protocadherin involved in regulating cancer cell motility ^62^. Finally, the promoter of *TP53* is ranked fifth, with overall near significance (q=0.28) and nominal significance for head and neck squamous cancer (p=0.043) as well as Chromophobe kidney cancer (p=0.006; **Supplementary Fig. 8d,e**). These mutations affect splice sites and thus post-transcriptional regulation.

The miR122 promoter region is third-ranked (q=0.22), with nominal significance in liver hepatocellular carcinoma (HCC; p=0.022). The miR122 region was originally detected as a driver candidate based on liver cancer indel mutations (q=0.043; **Supplementary Table 2 and** Figure 6a). The liver cancer mutations (n=5) from the exome set were also primarily INDELs (n=3). The exome mutations were generally centered around the precursor miRNA (pre-miRNA), though this is probably a consequence of it being included in the capture. In addition, skin-cancer mutations also overlap pre-miR122, though mostly lacking survival data (**Figure 6b**). Interestingly, low levels of miR-122 is associated with poor prognosis in hepatocellular cancer (HCC)^63,64^, where it has been discussed as a therapeutic target^65^.

By use of same sample miRNA profiles, we asked if the mutations in miR122 were associated with low miR122 expression levels (**Figure 6c**). This was generally the case, though the effect was only significant compared to normals (p=2.6x10^−7^) and not HCC cancers (p=0.13), which are generally down-regulated. We also asked if the protein-coding genes with 3’UTR miR122 target sites were significantly perturbed as a set. This was the case for a patient (A122) with a four-bp deletion that affects the 5’ end of miR122 (p=2.4x10^−9^; see Methods). In general a highly significant correlation between miR122 expression levels and target gene perturbation was observed in HCC samples (p=8.1x10^−28^). The patient with a four-bp deletion and the lowest miR122 expression level has the shortest overall survival of the five (**Figure 6d**).

## Discussion

Our two-stage procedure, ncDriver, identified non-coding elements with elevated conservation and cancer specificity of their mutations, which were further characterized by correlation with expression and survival to shortlist non-coding driver candidates. Importantly, the procedure is designed to be robust to variation in the mutation rate along the genome, as significance evaluation and candidate selection is based on surprising mutational properties, given sequence context, and not the overall rate. In addition to recovering known protein-coding drivers, it top-ranked known non-coding driver elements, such as promoters and enhancers of *TERT* and *PAX5*^*3,8,9,12*^. It also recalled a surprising intensity and distribution of mutations in *CTCF* binding sites of the cohesin complex^34^, which were found to correlate with high conservation and DNase I hypersensitivity.

Distinguishing non-coding driver elements shaped by recurrent positive selection from localised mutational mechanisms and technical artefacts is challenging. It may therefore be only a minority of the identified significant elements that are indeed true drivers, which stresses the importance of careful case-based analysis. To assist in the prioritization and shortlisting of non-coding driver candidates, we systematically evaluated the association of mutations in the identified elements with expression as well as patient overall survival using independent data sets. The expression correlation identified known drivers, an increased correlation at recurrently mutated TFBS sites, and pinpointed individual recurrently mutated candidate elements with strong mutation-to-expression correlations. Similarly, the survival analysis top-ranked known protein-coding and non-coding drivers, identified non-coding candidates where mutations associated significantly with decreased survival for individual cancer types, and supported miR-122 as a potential non-coding driver in hepatocellular liver cancer.

In general, few non-coding elements showed the same level of mutational significance as the known protein-coding drivers. The integration of multiple sources of evidence therefore becomes necessary for robust detection. We found the introduction of a cancer specificity test contributed both to the top ranking of known driver elements and the evidence underlying some novel candidates. Similarly, integration of both expression and patient survival data may provide further insight into the functional impact and driver potential of mutations^14^. With low recurrence and few mutations, we evaluated only pre-selected candidate elements that passed a mutational recurrence test and thereby retained power compared to a more inclusive screening approach.

Some driver mutations may only affect gene expression in early cancer stages and be undetectable by the expression analysis. On the other hand, passenger mutations could potentially affect expression without affecting cell survival. However, the much higher expression correlation signal among double-hit than single-hit mutations in *CTCF* binding sites is compatible with a selective enrichment for functional impact and hence presence of driver mutations. However, mutational mechanisms may also correlate with expression in some cases (see below)^22,23^.

Similarly, some driver mutations may affect cancer onset but not disease progression and overall survival. Even if the mutations do affect survival, the effect has to be relatively large to be detected with the current cohort sizes and the small numbers of mutated elements for individual cancer types.

On the other hand, mutational processes may lead to false positive driver candidates in some cases. Although the cancer specificity tests model the cancer-specific context-dependent mutation rates in each element type, highly localized and potentially uncharacterized mutational processes may inflate the false discovery rate. Specifically, somatic hypermutation in lymphomas appear to underlie the significance of several of the transcription-start-site proximal top-ranked elements.

Here a mutational mechanism may therefore explain overall mutational recurrence and cancer-type specificity – additional evidence is needed to support them as driver candidates. Nonetheless, some of these also exhibit an enrichment of mutations affecting highly conserved positions, including the intronic *PAX5* enhancer and the *DMD* promoter, suggesting that there may be an enrichment of driver mutations that affect function. The expression-correlation analysis also top-ranked known targets of somatic-hypermutation (*MYC* and *BCL6*; **Fig. 5**). However, correlation between somatic hypermutations and expression level as well as translocation of some genes to immunoglobulin enhancers can explain this signal more parsimoniously^12,22^.

Several of the identified non-coding driver candidates are associated with chromatin regulation, either through association to regulatory genes (e.g., *TOX3* intronic enhancer) or as binding sites for chromatin regulators (e.g., both *PAX5* enhancers and *CTCF* TFBS near *MAPRE3*). In addition, the full set of cohesin binding sites show elevated mutation rates^34^, though micro-environment specific mutational processes may potentially underlie most of these^66^. This could suggest a potential role of non-coding mutations in shaping chromatin structure during cancer development, which is supported by the recent finding of chromatin-affecting non-coding mutations that create a super-enhancer in lymphoblastic leukemia^42^. Systematic integration of sample-level chromatin data in large cancer genomics studies would help reveal the broader relationship between non-coding mutations and epigenomics, which may both be driven by mutational mechanisms and selection.

This study has identified elements with surprising mutational distributions and shortlisted a small number of non-coding driver candidates with mutations that associate with expression and patient survival across independent data sets. However, given the small number of mutated samples and the resulting lack of power, validation in large independent cohorts will be needed. The power to discover and validate non-coding driver elements will increase with larger sample sets and further integration of functional genomics and clinical data^67^, as will be provided by the next phases of TCGA and ICGC, providing a basis for biomarker discovery, precision medicine, and clinical use.

## Methods

### Pan-cancer whole-genome mutations and non-coding element annotations

Pan-cancer whole-genome mutations were extracted from a previous ICGC mutation signature study containing 3,382,751 single nucleotide variants (SNVs) from 507 samples of ten tumor types and 214,062 insertions and deletions (INDELs) from a subset of 265 samples of five tumor types (Fig. 1a; **Supplementary Table 1**)^7^. The INDELs were included by mapping them to their first (lowest) coordinate. All analysis is done in reference assembly GRCh37 (hg19) coordinates. INDELs were cleaned by removing those that overlap known common genetic polymorphisms identified in the thousand genomes project phase 3 version 5b (2013-05-02)^68^.

Annotations of protein-coding genes, lncRNAs, sncRNAs and pseudogenes were taken from GENCODE version 19, Basic set^17^. Only coding-sequence features were included for protein-coding genes. Promoter elements of size 1 kb and 4 kb were defined symmetrically around GENCODE transcription start sites (TSSs). Annotations of regulatory elements included DHSs, transcription factor binding site peaks (TFPs), TFBS motifs in peak regions (TPMs) and enhancers were taken from a previously compiled set^18^.

ENCODE blacklisted regions that are prone to read mapping errors were subtracted from all elements^24^. CRG low-mappability regions, where 100-mers do not map uniquely with up to two mismatches, were downloaded from the UCSC Genome Browser and subtracted^69^. Finally, hyper-mutated genomic segments containing GENCODE Immunoglobulin and T-cell receptor genes together with 10 kb flanking regions, combined when closer than 100kb, were also subtracted. All non-coding elements were subtracted coding sequence regions, to eliminate detection of potential protein-coding driver mutations in these.

The processed lists of 10,982,763 input elements consisted of 56,652 transcripts for 20,020 protein-coding genes, 17,886 transcripts for 13,611 lncRNA genes, 8,836 transcript for 6,948 sncRNA genes, 948 transcripts for 889 pseudogenes, 94,465 promoters of size 1 kb for 41,598 genes, 94,956 promoters of size 4 kb for 41,875 genes, 2,853,220 DHSs, 417,832 enhancers, 5,677,548 TFPs and 1,760,420 TPMs (Fig. 1c).

Mutations were mapped to elements using the intersectBed program of the BEDTools package (Quinlan and Hall 2010). To avoid large signal contributions from individual samples, no more than two randomly selected mutations were considered per sample in any individual element.

### Two-stage procedure for identifying non-coding elements with conserved and cancer specific mutations

A two-stage test procedure, named ncDriver, was developed to evaluate the significance of elevated conservation and cancer specificity of mutations in non-coding elements (Fig. 1d), which was applied to each combination of mutation type and element type (Fig. 1a,c). The first stage identified genomic elements with surprisingly many mutations (high recurrence) and the second assigned significance to each of these according to the element mutation properties in terms of cancer specificity and conservation. Importantly, the two stages are independent of each other, as the property tests are conditional on the number of mutations. Final significance evaluation and element selection was based only on the mutations properties, not their recurrence, to increase robustness against rate variation between samples and along the genome^1^. The first stage thus acts as a filtering step of elements considered for candidate selection. Details of the stages and involved tests are given below.

*Mutational recurrence test*. The recurrence test evaluated if the total number of mutations in an element was surprisingly high given its lengths and the background mutation rate for the given element type based on a binomial distribution. In case of overlapping elements, the most significant element was selected. P-values were corrected for multiple testing using the Benjamini and Hochberg procedure (BH)^70^ and only elements passing a 25% FDR threshold were passed on to the second stage.

In the second stage, three separate tests evaluated the cancer specificity and conservation of the mutations within each element. *1) Cancer-specificity test; 2) Local conservation test*: average conservation level of mutated positions compared to a local element-specific distribution and *3) Global conservation test*: average conservation-level of individual mutated positions compared to the genome-wide distribution for the element type.

*1) Cancer-specificity test*. For each element, the number of observed mutations in each cancer type was calculated. The expected number of mutations was also calculated for given element type and cancer type, grouped by mutation trinucleotide context to account for individual cancer type mutation signatures. We then asked if the distribution of observed mutations across cancer types within the element was surprising compared to the expected number of mutations using a Goodness-of-fit test with Monte Carlo simulation, (Fig. 1d.i). In the local and global conservation tests, we evaluated for each element if the mutations were biased toward highly conserved positions and thus potentially of high functional-impact. *2) Local conservation test*. In the local conservation test, the p-value of the mean phyloP conservation score^71^ across the observed mutations was evaluated in an empirical score distribution derived from 100,000 random samples with the same number of mutations and the same distribution of phyloP scores as the element in question (Fig. 1d.ii). *3) Global conservation test*. In the global conservation test, we applied the same sampling procedure to evaluate if mutations hit positions of surprising high conservation compared to the observed distribution across all elements of the given type (Fig. 1d.iii). Fisher’s method was used to combine the three individual p-values of the second stage to an overall significance measure. Again, p-values were corrected using BH and a 25% FDR threshold was applied to generate the final ranked candidate element lists.

### Driver recall in known cancer genes and an independent whole-genomes data set

Driver recall in known cancer genes were evaluated by the number of genes, associated with significant elements, that overlap genes in the COSMIC Gene Census database version 76^72^. Significance of observed enrichments were calculated using Fisher’s exact test for two-times-two contingency tables (**Supplementary Table 3**).

Recall of individual candidate driver elements was evaluated in an independent mutation data set from 505 whole-genomes with 14,720,466 SNVs and 2,543,085 INDELs^14^ (**Supplementary Fig. 1**). Using the list of 208 unique, non-overlapping and significant elements (48 protein-coding and 160 non-coding), we defined a *single elements* and a set containing *gene level elements* for recall testing using ncDriver (**Supplementary Fig. 1a**). The *single elements* set (n=208) simply consisted of all significant elements, whereas the *gene level elements* set (n=251,333) contained all elements sharing the same associated gene IDs (by nearest protein-coding gene for regulatory elements) as the individual significant elements. The *single elements* were analyzed as a single set, whereas the *gene level elements* were analyzed per element type, in both sets applying the ncDriver procedure to identify significantly recalled elements (**Supplementary Fig. 1b**). The significantly recalled elements were further analyzed for mutation correlation with patient survival as described in the Methods section ‘Two-stage procedure for identifying non-coding elements with conserved and cancer specific mutations’.

The observed number of recalled elements in the *single elements set* was evaluated by significance for each element type using monte carlo simulations (**Supplementary Fig. 1b**). The same number of elements as in the candidate set (n=208) were randomly drawn from the input element set, while the maintaining the relative distribution between element types. Each random element set was then subjected to ncDriver, the same procedure, which was used to detect the significant elements in the original data set. The p-value of the number of recalls for the original data set was evaluated as the fraction of random sets that led to the same (m) or a higher number of recalls (p=(m+1)/(1000+1))^73^; **Supplementary Table 4**). The ncDriver driver screen procedure is described in the Methods section ‘Two-stage non-coding driver detection’.

### Correlation of mutations in non-coding elements with gene expression

Exome mutations from 5,802 patient samples for 22 cancer types were downloaded from TCGA^16^. Somatic mutations with the PASS annotation were extracted and cleaned for genetic polymorphisms by subtracting variants from dbSNP version 138. A final set of 5,621,521 mutations was created, representing 2,726,008 INDELs and 2,895,513 SNVs. Mutations found in elements detected as significant by ncDriver were extracted and annotated with gene names (using gene name of nearest transcription start site for regulatory element) and sample ID for expression correlation analysis (Fig. 5a-f).

TCGA expression data for 7,382 cancers from 22 cancer types (ACC (n=79), BLCA (n=408), BRCA (n=1,097), CESC (n=305), COAD (n=286), DLBC (n=48), GBM (n=152), HNSC (n=520), KICH (n=66), KIRC (n=533), KIRP (n=290), LGG (n=516), LIHC (n=371), LUAD (n=515), LUSC (n=501), OV (n=262), PRAD (n=497), READ (n=94), SKCM (n=104), THCA (n=505), UCEC (n=176) and UCS (n=57)) was obtained using TCGA-Assembler^74^. Expression calls for all genes (n=20,525) were log2-transformed and z-score-normalized within each cancer type.

Expressions on the z-score scale were combined for all cancer types and Wilcoxon rank-sum test scores were calculated following addition of a rank robust small random value to break ties. In the rank-sum test procedure, all samples for which no mutations were observed were considered non-mutated. All samples were used in the expression correlation analysis, though only a subset (n=4,128) had paired exome DNAseq mutation calls. For all genes with mutations in a given element type, a combined p-value was calculated using Fisher’s method for combined p-values.

### Correlation of miR-122 target site and expression

In each of 266 TCGA liver samples, a gene expression fold change value was calculated by dividing with the gene median expression of the normal liver samples. For each sample, genes were ranked by the fold change value. We used the R package Regmex^75^ to calculate rank enrichment of miR-122 target sites in the 3’UTR sequences of the genes. The motif enrichment is a signed score corresponding in magnitude to the logarithm of the p-value for observing the enrichment given the sequences and their ranking. Negative values corresponds to observing the target more often in genes expressed higher than the median level. The motif enrichment score was correlated with the expression of miR-122 in the liver samples.

### Association of mutations in non-coding elements with patient survival

To further evaluate the driver potential of the identified significant elements, we correlated the mutation status with survival data. We downloaded clinical data from the TCGA data portal^76^ (2015-11-01) using the RTCGAToolbox R library^77^. For a given element, the difference in survival between mutated and non-mutated samples was tested per cancer type using Log-rank test on the Kaplan-Meier estimated survival curves^78^. We specifically tested a hypothesis that the presence of candidate mutations decreases the survival^55^. For this, we fitted cox proportional hazard models to robustly determine the effect direction (increase versus decrease in survival), and then halved the Log-rank test p-value, which is the goodness of fit test in its essence, if the effect was in expected direction. To avoid evaluating the hypothesis in underpowered cancer types, the tests were only performed when at least three mutations were present. Evidence was combined across cancer types using Fisher’s method.

## Data Availability

UCSC track hubs for significant elements and script codes for the ncDriver procedure can be obtained using the following URL: http://moma.ki.au.dk/ncDriver/.

## Acknowledgements

We thank the International Cancer Genome Consortium (ICGC) and The Cancer Genome Atlas (TCGA) for access to cancer genomics data sets. This work was supported by the Sapere Aude program from the Danish Councils for Independent Research, Medical Sciences (FSS), an Aarhus University Interdisciplinary Network Grant, and an Aarhus University interdisciplinary research centre grant (iSeq).

## Author contributions

J.S.P. initiated and led the analysis. H.H., M.M., N.A.S.-A., R.S., T.Ø., A.H., and J.S.P. contributed to analysis design. H.H. conducted mutational analysis, with assistance from M.J. and T.M.. N.A.S.-A., R.S., and M.K. conducted epigenomic and IGR impact analysis. M.M. conducted mutation to expression correlation analysis. MPS performed the survival correlation analysis. H.H., M.M., and J.S.P. wrote the paper with input from all other authors.

